# Selective attention modulates surface filling-in

**DOI:** 10.1101/221150

**Authors:** William J. Harrison, Alvin J. Ayeni, Peter J. Bex

## Abstract

**The visual system is required to compute objects from partial image structure so that figures can be segmented from their backgrounds. Although early clinical, behavioral, and modeling data suggested that such computations are performed pre-attentively, recent neurophysiological evidence suggests that surface filling-in is influenced by attention. In the present study we developed a variant of the classical Kanizsa illusory triangle to investigate whether voluntary attention modulates perceptual filling-in. Our figure consists of “pacmen” positioned at the tips of an illusory 6-point star and alternating in polarity such that two illusory triangles are implied to compete with one another within the figure. On each trial, observers were cued to attend to only one triangle, and then compared its lightness with a matching texture-defined triangle. We found that perceived lightness of the illusory shape depended on the polarity of pacmen framing the attended triangle, although the magnitude of this effect was weaker than when all inducers were of the same polarity. Our findings thus reveal that voluntary attention can influence lightness filling-in, and provide important data linking neurophysiological effects to phenomenology.**

## Introduction

The natural environment is cluttered with mutual occlusions among objects. Our ability to group common parts of goal-relevant objects while segmenting them from distracting objects is a critical visual function. Visual illusions provide powerful tools to probe the neural computations involved in such perceptual organisation. Since being popularised several decades ago ^1^, psychologists, neuroscientists, and philosophers have used Kanizsa figures to debate the mechanisms underlying the perception of occlusions, lightness and form. These figures give rise to a vivid percept of a shape emerging from sparse information, and thus demonstrate the visual system’s ability to interpolate structure from fragmented information, to perceive edges in the absence of luminance discontinuities, and to fill-in a shape’s surface properties ^1^.

Visual attention – focusing on some parts of an image – is known to modulate the perception of figure-ground relationships. Driver and Baylis ^2^ found that figure-ground organization of ambiguous figures was influenced by the region to which observers attended. Attended regions were interpreted as figures, whereas unattended regions were interpreted as ground. In addition to manipulating depth order, Tse ^3^ found that voluntary attention can influence surface filling-in. He developed a visual illusion that demonstrates that voluntary attention modulates the lightness of overlapping transparent surfaces. In one version of this illusion, three discs are positioned around a fixation point such that their borders overlap slightly. By covertly shifting attention to each disc in succession, the viewer can perceive the attended disc to be darker than the others, thus revealing that visual attention influences filling-in processes. In these cases, as well as others including Rubin’s classic face-vase figure or the Necker cube, visual attention modulates the appearance of surfaces that are defined physically. It remains unclear, however, whether visual attention modulates the visual system’s representation of illusory structure – i.e. structures that are generated by visual processing itself.

Psychophysical findings that visual attention influences figure-ground segmentation are consistent with the notion that visual attention acts on structure computed by border ownership cells. Qiu et al ^4^ investigated whether the responses of border-ownership cells in macaque area V2 are influenced by visual attention. Border-ownership cells signal which side is an object versus background for a given border. After determining the side preference of a sample of border-ownership cells, they displayed overlapping figures such that a border shared between two objects was positioned within the cells’ receptive fields. They found that some border-ownership cells signal figure-ground relationships in the absence of visual attention, but that the activity of other cells is modulated by visual attention. Qiu et al suggested that visual attention can influence figure-ground perception by modulating the gain of border-ownership cells, biasing perception more toward one percept over a competing one. Poort et al ^5^ further investigated the modulatory effects of attention on the responses of V1 and V4 neurons to object borders and surfaces. They found that, whereas the response of neurons to an object’s borders were relatively unaffected by attentional allocation, filling-in of an object’s surface was modulated by attention.

In contrast to the above studies suggesting an influence of attention over perceptual organization, other investigations suggested perception of illusory Kanizsa figures is pre-attentive. Mattingley et al ^6^ investigated whether Kanizsa figures were perceived by a stroke patient who experiences “extinction”, a phenomenal loss of awareness of contralesional stimuli when presented concurrently with ipsilesional stimuli. Despite gross lapses in attentional allocation to the contralesional space, this patient was nonetheless able to perceive Kanizsa figures whose illusory borders extended across the visual meridian. This finding suggests that filling-in is a pre-attentive process. Davis and Driver ^7^ drew a similar conclusion after having healthy participants perform a visual search task involving illusory figures. They found that the time taken to find an illusory figure in a display did not increase with additional search items, a hallmark of pre-attentive processing (but see reference ^8^). The role of visual attention in computing illusory objects thus remains somewhat contentious.

In the present study, we designed a multi-stable illusory figure to investigate whether observers’ visual attention can determine perceptual filling-in of illusory figures. Although an object’s boundary is typically defined by luminance, color, or texture contrast, borders can be perceived where no physical difference exists when spatially discontiguous visual features are interpolated to create illusory contours. The illusory edges of the sort produced by Kanizsa figures composed of isolated ‘pacman’ inducers elicit contour responses in V2 neurons ^9^ and produce a vivid percept of a shape. Of particular importance to the present study, the surface of Kanizsa figures are filled-in, resulting in an apparent surface lightness of greater contrast than its immediate surrounds ^1,10^. Kanizsa figures thus offer important insights into the mechanisms underlying perceptual organization by providing minimal conditions under which multiple phenomena arise.

Our new figure is composed of two spatially overlapping Kanizsa triangles (Fig. 1; Gaussian blurred to strengthen the effect, see also Fig. 2). On first inspection, the reader may perceive a grey star occluding six black and white discs. However, the black and white inducers are arranged so as to imply competing geometric forms. Whereas the black inducers imply an inverted triangle, the white inducers imply an upright triangle. These cues are equally physically salient so that, in the absence of attention, perceptual organization may be a random draw of any possible interpretation (e.g. a star, an upright triangle on top, or an inverted triangle on top). Thus, with a single image, we can manipulate observers’ attention and test whether there is a corresponding systematic change in filling-in.

**Figure 1.**
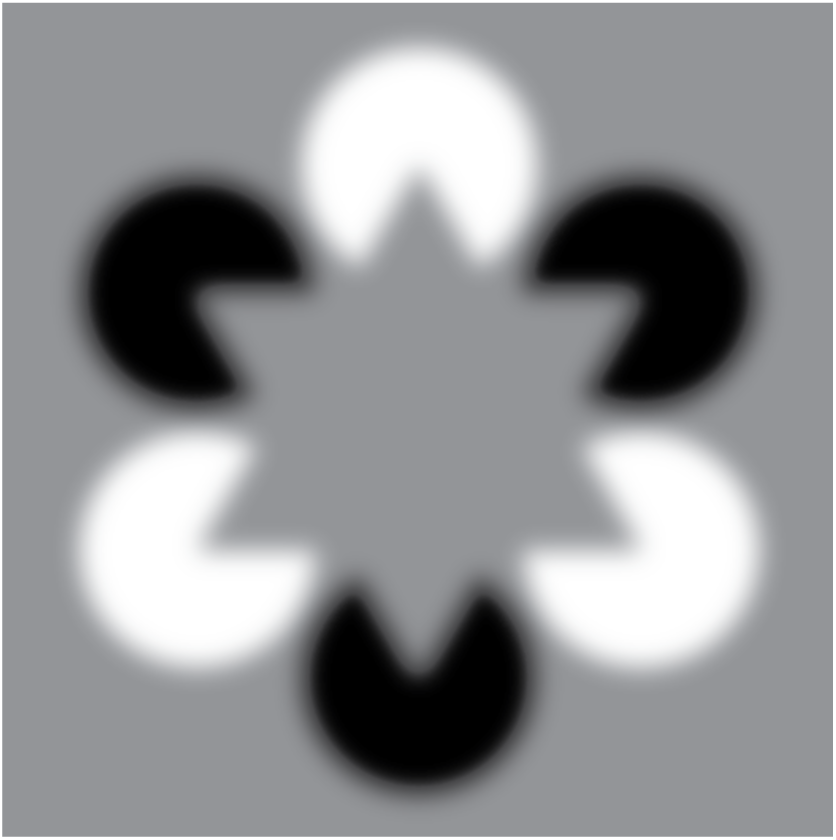
A novel Kanizsa figure to test attention-contingent filling-in. When attending to the black inducers, a downward-pointing triangle emerge as the top-most surface, and this attention-contingent illusory triangle may appear lighter than its background. When attending to the white inducers, a darker, upward pointing triangle may appear forming the top-most object, and this may now appear darker than its background.

**Figure 2.**
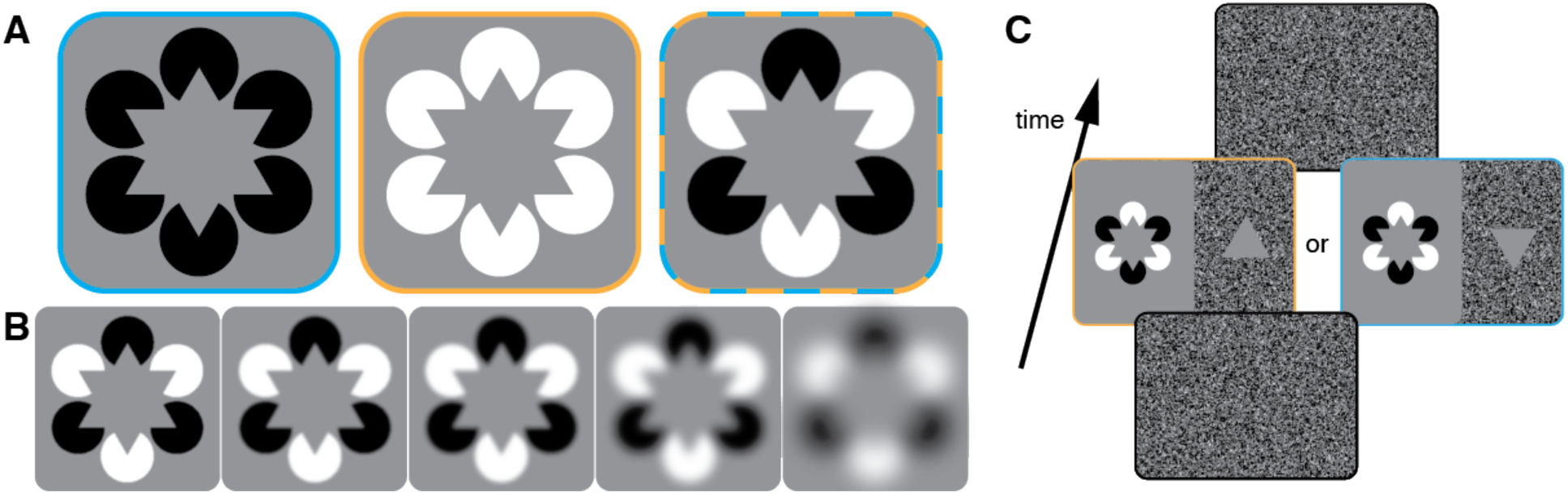
Examples of all experimental stimuli and procedure. A) In the classical Kanizsa figures ^1^, inducers within a figure were equal in luminance. In our experiment, we thus included “homogeneous inducer” conditions in which the inducers were all black (blue outline) or all white (orange outline). In these versions, the illusory hexagrams appear lighter and darker than their backgrounds, respectively. These conditions provided a relative baseline against which we could compare perception in our new “mixed inducer” figure (blue and orange outline). B) Illustrative examples of blur levels. Across conditions, we applied various Gaussian blur levels (s=0.1^°^ to 30° in log steps) to the inducers and to the white noise background (see (C) and Methods) in order to vary uncertainty about the illusory edges of the Kanizsa figure. Shown from left to right are the least to most blurred mixed inducer conditions. Note that in the most blurred condition, the sharp edges of the inducers were abolished. C) Two example mixed inducer trials. An observer reported whether the texture-defined matching triangle (shown here on the right half of the middle displays) was lighter or darker than the illusory triangle with the same orientation. In this example, the matching triangle implicitly cued the observer to attend to the illusory triangle in white inducers in the left stream, and to the illusory triangle in black inducers in the right stream. See Methods for details.

We designed the stimulus in Figure 1 to control for the known lightness effect observed in the classic Kanizsa figure. In Figure 2A we show our novel stimulus alongside two other variants of the classic figure in which the inducers are all of the same polarity. When inducers are homogenous in their luminance, any influence of attention on the illusory surface could minimise or amplify the baseline illusion, and such an effect could be explained simply by an interaction of attention with the low-level stimulus properties. In contrast, we designed the mixed-inducer condition such that the perceptual outcome cannot be predicted by the low-level stimulus properties alone -- depth order and filling-in must be determined either stochastically, or according to an observer’s selective attention. It was also critical for our stimulus to imply two spatially overlapping figures to test the hypothesis that depth order is determined prior to the stage at which visual attention operates e.g. ^6,11^. Had we presented two spatially non-overlapping illusory figures, one defined by black inducers and the other defined by white inducers, any modulatory effect of attention on filling-in could be attributed to pre-computed structure (e.g. compare the apparent surface lightness of the homogenous-inducer stimuli of Fig. 2A). Therefore, the spatially overlapping implied triangles of our mixed-inducer condition allows us to assess if selective attention can modulate depth order of illusory surface properties inferred by the visual system.

## Results

We investigated whether filling-in of an illusory surface can be modulated by voluntary endogenous attention. A typical trial sequence of our psychophysical task is shown in Figure 2C. Observers (n = 15) reported the apparent lightness of only the upward oriented illusory triangle or only the downward oriented illusory triangle by comparing it with a luminance-defined matching triangle presented on a background of white noise. To draw observers’ attention to one of the two illusory triangles, they were instructed to judge only the figure that matched the orientation of the luminance-defined triangle. We thus directed observers to attend to an illusory triangle defined by white (or black) inducers endogenously without making reference to the colour of the inducers. We refer to this condition as the “mixed inducer” condition. For comparison, we also included conditions in which all inducers were black or white (“homogenous inducer” condition; Fig. 2A). Whereas the critical mixed inducer condition allows us to investigate clearly the influence of voluntary attention on perceptual filling-in, the homogenous inducer condition provides a baseline in which conflict between competing structural cues is reduced. Finally, we further tested whether filling-in was affected by certainty of edge location by applying varying levels of blur to the Kanzisa figure and matching triangle background (Fig. 2B). The inducer condition, orientation of the illusory figure, matching triangle, level of blur, and display side of the matching triangle were all counter-balanced across trials (see Methods for all experimental details).

We defined perceived lightness as a point of subjective equality (PSE), the mid-point of the psychometric function fit to the proportion of “lighter” responses as a function of the contrast of the matching triangle. Psychometric functions fit to data collapsed across participants and blur condition are shown in Figure 3. These fits were calculated without the most blurred condition in which the results were different than the other conditions (see Fig. 4A).

**Figure 3.**
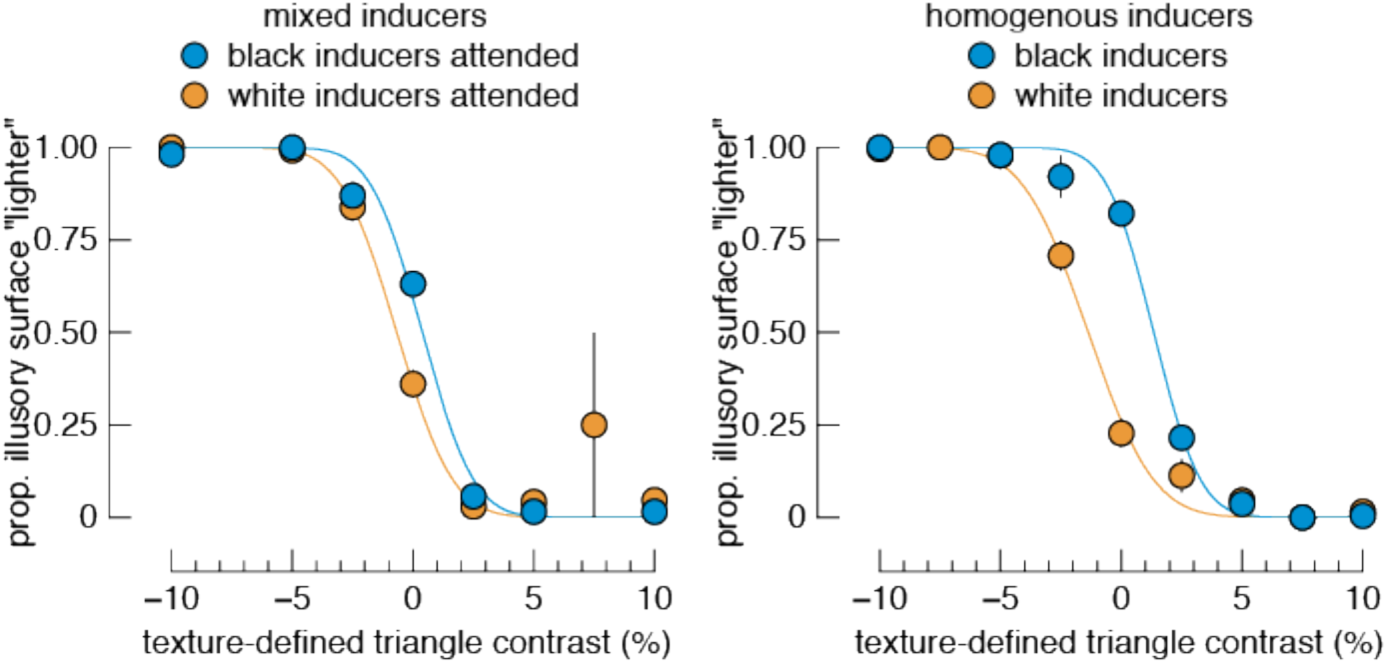
Mean performance and psychometric functions. Data show the mean proportion of times observers reported the illusory surface as “lighter” than the texture-defined surface, as a function of the texture-defined triangle contrast. These data are averaged over all blur conditions except the most blurred condition, which was omitted due to its obscuring of the main effects (see Fig. 4A). Despite being shown the same stimulus in the “mixed inducer” condition, there is a difference in curves according to the attended inducer polarity (left panel). On average the effect is weaker than in the “homogenous inducer” condition (right panel). Note that for the main analyses presented in the Results, data were fit separately for each participant and each condition. Error bars show one standard error.

**Figure 4.**
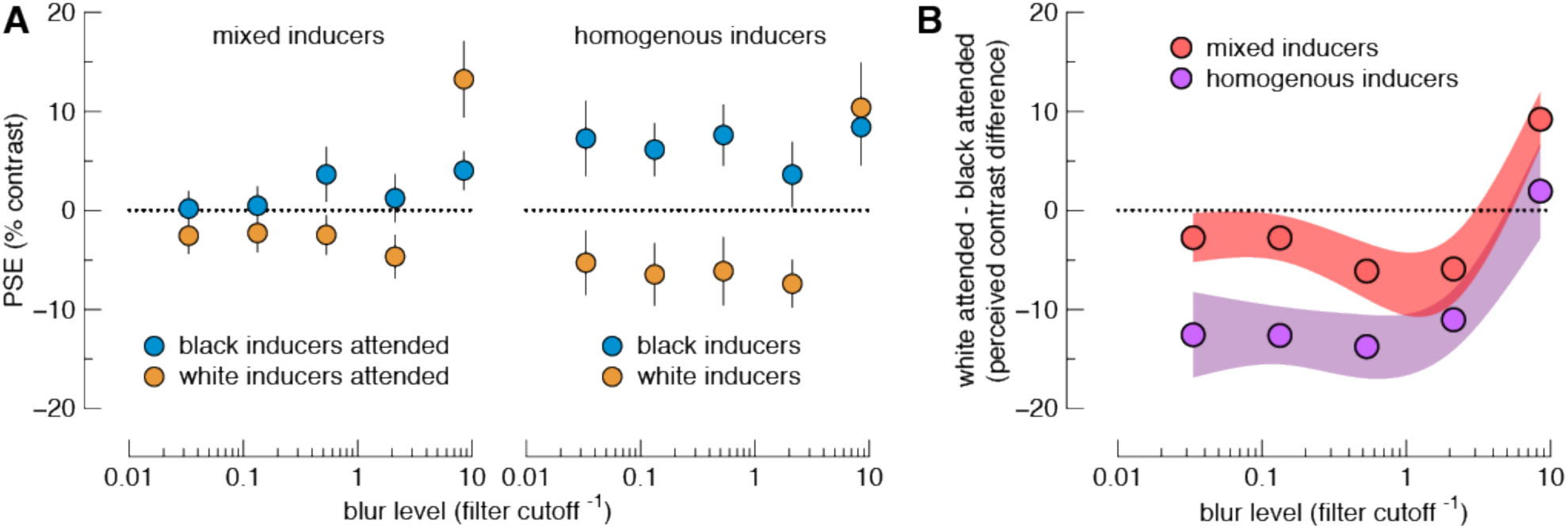
Extended psychophysical data. A) PSEs from all tested conditions. Data show the point of subjective equality (PSE) for each condition over a range of blur levels (see Fig. 3). Positive and negative values on the ordinate indicate the attended surface was perceived as lighter or darker, respectively, than its actual luminance. Higher values on the abscissa indicate greater levels of blur. B) To facilitate comparison across all conditions, we plot the difference in PSEs between attend white inducer and attend black inducer conditions (ordinate). Data points thus show the perceived lightness difference between illusory surfaces embedded in white versus black inducer elements over a range of blur levels, with either mixed or homogeneous inducers. Negative values on the ordinate indicate that an attended surface in white inducers was perceived as darker than an attended surface in black inducers. Error bars in (A) show one standard error. Shaded regions in (B) show 95% confidence intervals. N = 15.

PSEs for all conditions, averaged across participants, are shown in the right panel of Figure 4A. For all but the most blurred condition of the mixed inducer condition, illusory triangles defined by black inducers were perceived as lighter than illusory triangles defined by white inducers. This effect appears to be weakest for the two least blurred conditions. We verified these observations with Bayesian paired samples t-tests, comparing PSEs across inducer polarity within each blur level. There was strong evidence of a difference for the three most blurred conditions, but equivocal evidence of a difference of PSEs within the two least blurred conditions (BF10 in order of least to most blurred = 0.821, 1.627, 66.64, 10.585, and 80.393). Therefore, despite being shown the same image (within each blur condition), observers’ perceptual reports depend on their attentional goals.

We included a homogenous inducer condition in which all inducers were of the same polarity in each trial. This condition provides little ambiguity over the interpretation of the figure, but we nonetheless instructed observers to attend to only the shape cued by the texture-defined triangle. The homogenous inducer condition thus gives an indication of the upper-bound of the difference in PSEs between black vs white inducer conditions. Data from this condition are shown in the right panel of Figure 4A, and reveal a stronger effect of inducer polarity than the mixed inducer condition, with equal strength across all but the most blurred condition. Bayesian paired samples t-tests revealed strong evidence of a difference in PSE between the inducer polarities for the four least blurred conditions, but evidence against a difference for the most blurred condition (BF_10_ in order of least to most blurred = 28.919, 427.841, 297.566, 105.704, and 0.304).

The strength of the effect of attention on PSEs is further summarized in Figure 4B. In this figure we show the difference between attending to the figure in black inducers vs attending to the figure in white inducers for all conditions. To formally test for whether there was a difference between mixed vs homogenous inducer conditions, we submitted these difference scores to a 5 x 2 repeated measures Bayesian ANOVA, with factors blur level (5 levels of blur), and inducer type (2 levels: mixed vs homogenous). The best model included the two main effects (blur level + inducer type: BF_10_ = 5.264e+17). There was strong evidence against an interaction between factors (BF_10_ = 0.116).

Finally, we found weak to moderate evidence *against* a difference in the slope parameter of the psychometric functions for all black vs white inducer comparisons in Figure 4A (all BF_10_ < 0.5; min = 0.262; max = 0.444).

## Discussion

We asked observers to allocate their attention to varying elements of a novel illusory figure and report the perceived lightness of the surface of the center of the figure. By presenting the same figure on each trial and only manipulating observers’ goals, we controlled physical aspects of the stimulus and found that observers’ reports of perceived lightness were contingent on to which illusory figure they attended. We do not think observers selectively reported the apparent lightness of only some parts of a pre-attentively perceived image in the mixed inducer condition for three reasons. First, the influence of selective attention over depth order and apparent surface lightness can be experienced first-hand by the reader by inspecting Figure 1. Second, we had observers verbalise their reports during an initial practice session (see Methods), and they frequently reported that the attended triangle appeared as the top-most surface. Third, Harrison and Rideaux have reported that selectively attending to only some inducers of the mixed-inducer stimulus results in the perception of illusory edges between those inducers ^12^, providing converging evidence that the depth ordering of the stimulus depends on observers’ attention control. We thus infer that apparent surface lightness of our novel illusory figure can be modulated by attention.

We found that the strength of filling-in was weaker in the mixed-inducer condition than the homogenous inducer condition (Fig. 4B). There could be several reasons for this. First, there is a larger signal implied by the homogenous inducer condition, since all inducers share the same polarity. Second, lapses in attention on any given trial will have more impact on the mixed-inducer condition in which the opposite polarity inducers imply competing surface arrangements. Third, the modulatory effects of attention on filling-in in our mixed-inducer condition may simply be weaker than in cases in which attentional selection plays no role, which may have been the case in the homogenous inducer condition if observers perceived a star. Addressing this issue requires further investigation. Nonetheless, our finding of a difference in perceived lightness when black versus white inducers were attended in the same image (mixed inducer condition) reveals that illusory lightness is contingent on observers’ attentional goals. For all but the most blurred conditions, we found filling-in was not substantially moderated by the amount of spatial blur added to the illusory figures induced by homogenous pacmen. This finding is consistent with the idea that filling-in involves broadly tuned spatial filters that are insensitive to high spatial frequencies. We did, however, find a reversal of the direction of filling-in for the most extreme blur (rightmost point in left panel of Fig. 4A). We speculate that the explanation for this is straightforward: because the inducers were so blurred that the implied tips of the triangles, and thus the illusory figure, were abolished. In this case, filling-in would be consistent with the contrast of the inducers, rather than opposite to them. It is also worth noting that observers have a large bias to report the inner area of the figure as lighter in their responses for the most blurred homogenous inducer condition. This observation is not predicted by current accounts of filling in but we speculate that it could be related to the blackshot mechanism ^13^ that amplifies the relative salience of the dark inducers.

It has been shown that visual attention can affect perception over short timescales in multiple ways, such as by increasing apparent contrast or spatial frequency ^14^. Any such effect in the present study requires an initial grouping stage, such that changes in contrast were selectively applied to only some features of the image. Selectively attending to only the inducers surrounding the cued target shape could cause illusory spreading of the apparent surface lightness. Whereas previous studies have shown that visual attention can influence depth order ^2^ or even surface lightness of semi-transparent discs ^3^, the case presented here is unique in that it shows observers’ task instructions alter the visual system’s inference, creating one of multiple competing *illusory* surface appearances and arrangements.

Recent evidence suggests that cells in V2 play an important role in this process ^4,11,15,16^. V2 cells are selective for an object’s borders and the relative position of the object. A network of such “figure-ownership” cells may be supported by grouping cells, thus encoding figure-ground structure essential for visual cognition. Indeed, recent work has shown that when visual attention is allocated to a region of space from which a grouping cell is currently receiving structured input, figure-ground segmentation is enhanced by feedback from the grouping cell to its connected figure-ownership cells ^4,11,15^. This model may account for our results. Attending to the upright (or inverted) triangle would result in the grouping cell enhancing the structured input to only the corresponding shape, biasing the processing of form and systematically altering perceptual organization. Our results are also consistent with recent neuro-imaging research in humans using electroencephalogram, in which top-down attention modulates early neural activity associated with illusory contour formation ^17^. Our study provides important data that supports the notion that such top-down signals modulate perceptual phenomenology.

Our study thus helps to bridge a gap between psychophysical and neurophysiological investigations of figure-ground segmentation. Consistent with the activity of V4 neurons found by Poort et al ^5^, we found that perceptual filling-in of an object’s surface is modulated by attention. However, unlike the conclusion of Poort et al that attention does not modulate the responses to the object’s borders, our preliminary data using a similar figure to the present study suggests border computations also depend on voluntary attention ^12^.

## Acknowledgments

The authors thank Dr Emily Wiecek for her constant critical feedback, without which we would not have developed our methods fully. We also thank Guido Maiello for providing feedback on earlier versions of the manuscript. This work was supported by NIH grant R01EY021553 (PJB) and the National Health and Medical Research Council of Australia APP1091257 (WJH).

## Author contributions

W.J.H. and P.J.B. designed the Kanizsa stimuli and psychophysical procedure. A.J.A. ran the experiment, and W.J.H. analyzed the data. W.J.H. and P.J.B. wrote the manuscript. All authors discussed the results and commented on the manuscript.

## Data availability

The data that support the findings of this study are available from the corresponding author upon request.

The authors declare no competing financial interests.

## Methods

### Participants

15 observers completed the psychophysical experiment, including the authors. All were naïve to the purposes of the experiment, except for authors W.J.H. and P.J.B. This experiment was approved by the Northeastern University Institutional Review Board, and methods were carried out in accordance with the relevant guidelines. All participants were fully informed as to the requirements of participation prior to providing their consent.

### Stimuli and procedure

Stimuli were programmed with the Psychophysics Toolbox ^18^,^19^ in MATLAB (MathWorks), and were displayed on a CRT (40 x 30 cm, 1024 x 768 pixels; 85 Hz refresh). Six inducer “pacmen” (diameter = 3^°^ visual angle; jaw angle = 60^°^) were arranged at the corners of an imaginary hexagram with a uniform gray background (luminance = 20 cd/m^2^). From trial to trial, the inducers could be all white or all black (homogenous condition), or mixed (see Fig. 2). Maximum luminance of the white inducers was 40 cd/m^2^, minimum luminance of the black inducers was 0 cd/m^2^. Blur conditions were achieved by applying a Gaussian spatial filter with standard deviation of 0.1, 0.5, 1.9, 7.5, or 30.1° (Fig. 2). The matching triangle was an equilateral triangle (edge length = 6°) whose luminance varied according to an observer’s responses (see below). The matching triangle was presented on a background of white noise (mean luminance = 20 cd/m^2^), to which the same spatial filter was applied as for the Kanizsa figure. The apex of the matching triangle pointed upward or downward, depending on the condition (see below).

Observers sat in a dimly lit room approximately 58 cm from the display with a keyboard in their lap. Prior to each trial, visual noise masked the entire display for 500 ms. The Kanizsa figure was then presented on one half of the monitor and the matching triangle was presented on the other half (Fig. 2C). The side of the Kanizsa figure (and therefore the matching triangle) was selected randomly on each trial. Observers were instructed to press the left arrow key if the matching triangle was darker than the corresponding illusory triangle, or the right arrow key if it was lighter. The stimulus remained on the screen until a response was made. Responses were not time-limited and reaction times were not recorded: observers were instructed to not respond until they attended to the cued triangle.

To estimate PSEs from all 20 conditions in a single testing session, we used a 2-down 2-up staircase procedure that varied the luminance of the matching triangle according to an observer’s response series for each condition. Although this staircase converges on an observer’s point of subjective equality (PSE) – the luminance at which the matching triangle appeared equal to the attended illusory triangle – we nonetheless fit psychometric functions to constrain the PSE estimate by all observations (see *Analyses* section below). The staircase started randomly between ±20% contrast to ensure fitted psychometric functions were constrained. The initial step size was 10%, and halved after each reversal. A separate staircase was run for each of the 20 conditions, with all conditions interleaved. A staircase for a given condition ended after seven reversals, after which the condition was removed from the trial sequence.

For each of the five blur conditions (Fig. 2B), the inducers and matching triangle were arranged into four conditions: homogenous black inducers, homogenous white inducers, mixed inducers with the matching triangle cueing the white inducers, mixed inducers with the matching triangle cueing the black inducers. In all conditions, the orientation of the matching triangle was randomized to match the appropriate target illusory figure. Thus, with the five blur conditions and four inducer conditions, each observer was tested on a total of 20 conditions. Conditions were presented in random order until at least seven button press reversals were registered for every condition.

Prior to testing, observers were introduced to the classic Kanizsa figure, and then our modified mixed-inducer figure (Fig. 1). The experimenter then explained the task using static images that exemplified each of the inducer and blur conditions. When describing that the matching triangle cued the observer to which of the two possible illusory triangles to respond, the experimenter never referred to the inducer color, but only to the upward or downward pointing illusory triangle. The participants completed about 20 trials of practice and verbalized their responses in order for the experimenter to check they understood the task. Observers were encouraged to actively look at the matching triangle and illusory triangle, and to take their time with each decision. Training and all conditions were tested in a single session lasting approximately 30 minutes.

### Analyses

Cumulative Gaussian functions were fit to the proportion of “lighter” responses at each luminance level of the matching triangle for each of the twenty conditions. The PSE was the luminance that produced 50% lighter responses, converted to Weber contrast:

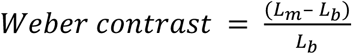

where *L_m_* is the luminance of the matching triangle producing 50% lighter responses for a given condition, and *L_b_* is the mean luminance of the background. This value was converted to a percent by multiplying by 100. Bayesian analyses were performed in JASP ^20^. In Figure 4B, we subtracted the mean PSE when the black inducers were cued from the mean PSE when the white inducers were cued for each blur condition.

## References

1. Kanizsa, G. Subjective contours. Sci. Am. 234, 48–52 (1976).

2. Driver, J. & Baylis, G. C. Edge-assignment and figure-ground segmentation in short-term visual matching. Cognit. Psychol. 31, 248–306 (1996).

3. Tse, P. U. Voluntary attention modulates the brightness of overlapping transparent surfaces. Vision Res. 45, 1095–1098 (2005).

4. Qiu, F. T., Sugihara, T. & von der Heydt, R. Figure-ground mechanisms provide structure for selective attention. Nat. Neurosci. 10, 1492–1499 (2007).

5. Poort, J. et al. The role of attention in figure-ground segregation in areas V1 and V4 of the visual cortex. Neuron 75, 143–156 (2012).

6. Mattingley, J. B., Davis, G. & Driver, J. Preattentive filling-in of visual surfaces in parietal extinction. Science 275, 671–674 (1997).

7. Davis, G. & Driver, J. Parallel detection of Kanizsa subjective figures in the human visual system. Nature 371, 791–793 (1994).

8. Gurnsey, R., Poirier, F. J. & Gascon, E. There is no evidence that Kanizsa-type subjective contours can be detected in parallel. Perception 25, 861–874 (1996).

9. von der Heydt, R., Peterhans, E. & Baumgartner, G. Illusory contours and cortical neuron responses. Science 224, 1260–1262 (1984).

10. Rieger, J. & Gegenfurtner, K. R. Contrast sensitivity and appearance in briefly presented illusory figures. Spat. Vis. 12, 329–344 (1999).

11. Craft, E., Schütze, H., Niebur, E. & von der Heydt, R. A neural model of figure-ground organization. J. Neurophysiol. 97, 4310–4326 (2007).

12. Harrison, W. J. & Rideaux, R. Voluntary control of illusory contour formation. (bioRxiv, 2017). doi:10.1101/219279 1.

13. Chubb, C., Landy, M. S. & Econopouly, J. A visual mechanism tuned to black. Vision Res. 44, 3223–3232 (2004).

14. Carrasco, M. Visual attention: the past 25 years. Vision Res. 51, 1484–1525 (2011).

15. Mihalas, S., Dong, Y., von der Heydt, R. & Niebur, E. Mechanisms of perceptual organization provide auto-zoom and auto-localization for attention to objects. Proc. Natl. Acad. Sci. U. S. A. 108, 7583–7588 (2011).

16. Zhou, H., Friedman, H. S. & von der Heydt, R. Coding of border ownership in monkey visual cortex. J. Neurosci. 20, 6594–6611 (2000).

17. Wu, X., Zhou, L., Qian, C., Gan, L. & Zhang, D. Attentional modulations of the early and later stages of the neural processing of visual completion. Sci. Rep. 5, 8346 (2015).

18. Pelli, D. G. The VideoToolbox software for visual psychophysics: Transforming numbers into movies. Spat. Vis. 10, 437–442 (1997).

19. Brainard, D. H. The Psychophysics Toolbox. Spat. Vis. 10, 433–436 (1997).

20. JASP Team. JASP (Version 0.8.5)[Computer software]. (2018).

